# The relative speed of the glucorticoid stress response varies independently of scope and is predicted by environmental variability and longevity across birds

**DOI:** 10.1101/2021.10.18.464833

**Authors:** Conor C. Taff, John C. Wingfield, Maren N. Vitousek

## Abstract

The acute glucocorticoid response is a key mediator of the coordinated vertebrate response to unpredictable challenges. Rapid glucocorticoid increases initiate changes that allow animals to cope with stressors. The scope of the glucocorticoid response — defined here as the absolute increase in glucocorticoids — is associated with individual differences in performance and varies across species with environment and life history. In addition to varying in scope, responses can differ enormously in speed; however, relatively little is known about whether speed and absolute glucocorticoid levels covary, how selection shapes speed, or what aspects of speed are important. We used corticosterone samples collected at 5 time points from 1,750 individuals of 60 species of birds to ask i) how the speed and scope of the glucocorticoid response covary and ii) whether variation in absolute or relative speed is predicted by environmental context or life history. As predicted by a recent model, faster absolute glucocorticoid responses were strongly associated with a larger scope. Despite this covariation, the relative speed of the glucocorticoid response (standardized within species) varied independently of absolute scope, suggesting that selection could operate on both features independently. Species with faster relative glucocorticoid responses lived in locations with more variable temperature and had shorter lifespans. Our results suggest that rapid changes associated with the speed of the glucocorticoid response, such as those occurring through non-genomic receptors, might be an important determinant of coping ability and we emphasize the need for studies designed to measure speed independently of absolute glucocorticoid levels.

## INTRODUCTION

Wild animals often encounter unpredictable and rapidly changing environmental conditions. For vertebrates, the glucocorticoid (GC) mediated stress response plays a primary role in coordinating phenotypic changes that allow animals to persist in challenging conditions (Sapolsky et al., 2000; Wingfield et al., 1998). Decades of evidence now demonstrate that rapid changes in GC hormones can alter a variety of downstream traits including metabolism, behavior, gene expression, and physiology in ways that promote the avoidance or tolerance of stressors (Dallman, 2005; Datson et al., 2008; Sapolsky et al., 2000; Wingfield et al., 1998).

While the basic structure of the GC response system is highly conserved (Romero and Gormally, 2019), individuals and species differ enormously in their absolute levels of circulating GCs under baseline and stress-induced conditions and in their regulation of GC levels (Romero and Gormally, 2019; Vitousek et al., 2019). Growing evidence suggests that observed differences in absolute GC levels among species reflect adaptation resulting from selection based on environmental context and life history (Bonier et al., 2009; Breuner et al., 2008; Cockrem, 2013; Schoenle et al., 2018; Vitousek et al., 2019; Williams, 2008). However, in addition to varying in the scope of the GC response, individuals and species may vary in the speed of response (see definitions in Box 1). In contrast to absolute levels, relatively little is known about how selection shapes the speed of GC responses.

The speed of the GC response might be an important target of selection if it determines how quickly individuals can match their phenotype to changing conditions (Luttbeg et al., 2021; Taff and Vitousek, 2016). Because the acute stress response is a multi-component system that includes a variety of downstream changes (Sapolsky et al., 2000), there will necessarily be a lag between the perception of any stressor and the production of the full stress-induced phenotype. Thus, a faster GC response should allow animals to more quickly match their phenotype with the prevailing environmental conditions (Taff and Vitousek, 2016). At the same time, responding faster might incur costs that could be avoided with a slower response, because prolonged or chronic elevation of GC levels can result in a variety of well known costs (Korte et al., 2005). Responding more slowly might allow animals to calibrate their response as additional information about a stressor is accumulated.

Disentangling the speed and scope of GC responses is challenging for several reasons. First, because the same physiological systems are involved in the speed and scope of the GC response, there are likely to be mechanistic links that create covariation between different attributes even when selection acts on only a single feature. For example, variation in FKBP5 expression could simultaneously alter the speed and magnitude of response (Zimmer et al., 2020a). Second, selection may favor the coupling of particular speed and scope combinations even when there is no intrinsic mechanistic link. For example, Luttbeg et al (2021) recently used optimality modeling of the speed of acute stress responses to show that altering GC regulation rate changes the optimal baseline and stress-induced GC levels under a variety of conditions. Finally, from a purely logistical perspective, separately measuring the speed and scope of stress responses is technically challenging (Taff, 2021). In simulations across a range of conditions, the most commonly used study designs have much higher power to detect variation in scope even when substantial variation in speed exists (Taff, 2021). Moreover, because variation in speed and scope can both contribute to differences in absolute GCs for a given sample (Box 1), variation in speed can be interpreted as variation in scope when samples are collected at standardized times (Figure 1, Taff, 2021).

**Figure 1:**
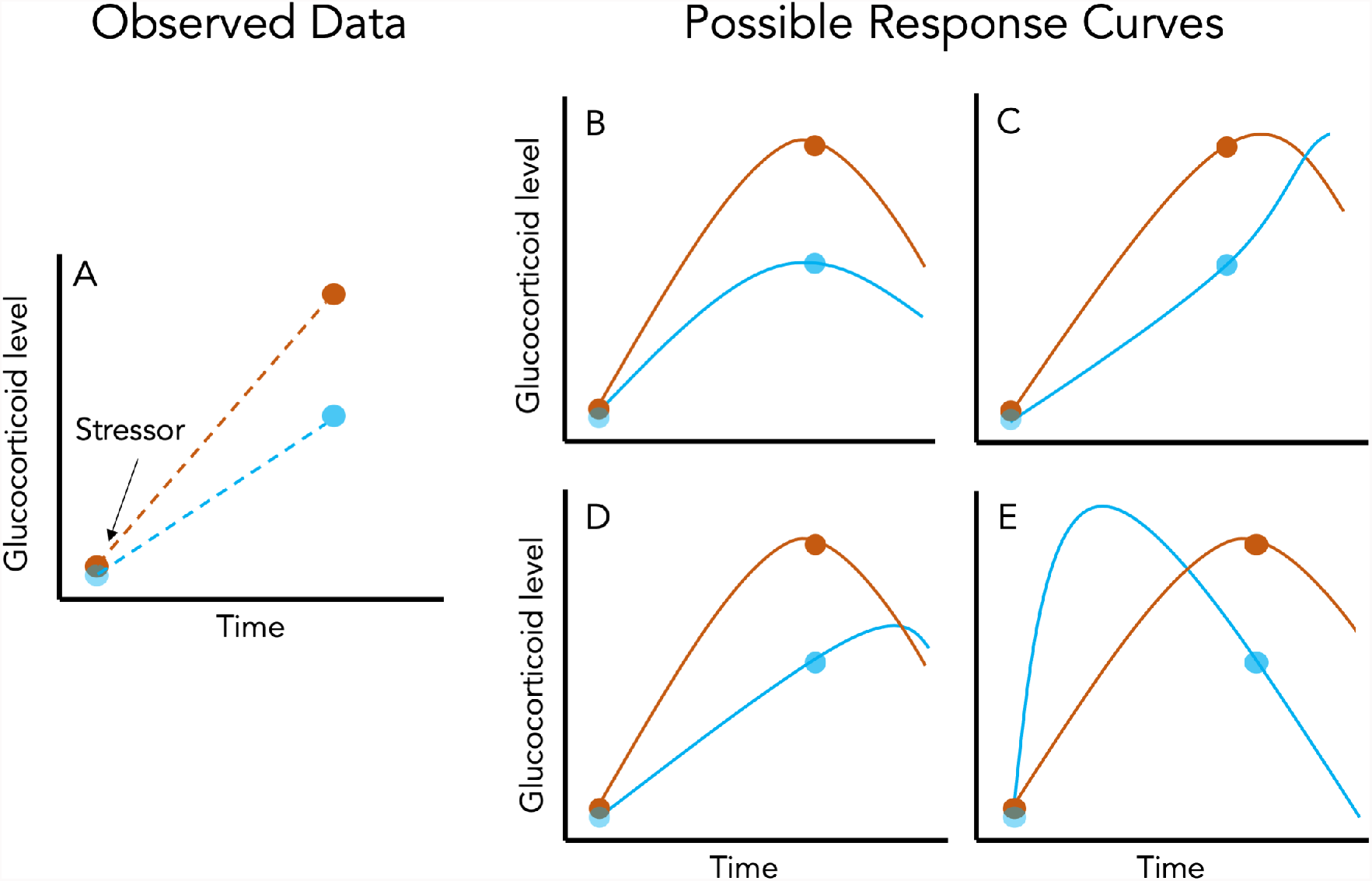
Panel A illustrates the data available when only a baseline and stress-induced sample are collected (points) from each individual (different colors) during a standardized stressor protocol. Dashed lines are the inferred increase in glucocorticoids in the interval between sampling. Panel B-E illustrate four different patterns of true acute responses (solid lines) that could produce the same observed data from Panel A. Individuals might differ only in the scope of their response with no variation in speed (B) or they might differ in the speed of the response without differing in scope (C). Alternatively, individuals might differ in both speed and scope (D and E). Depending on the nature of variation in both attributes and the timing of sampling, data collected at two time points might still capture variation in speed and scope (D) or it might entirely misrepresent variation in speed and scope (E). Increasing the number of sampled time points per individual will result in more constrained response curves, but the consequences of different sampling schemes are rarely considered.

Given the challenge of measuring the speed of GC responses, it is not surprising that there is much more empirical evidence suggesting the importance of variation in scope (Schoenle et al., 2018; e.g., Vitousek et al., 2019). However, there are also suggestions in the literature that variation in speed might differ in important ways between individuals in some situations. For example, wild great tits (*Parus major*) that were more cautious in a behavioral assay also had a faster increase in corticosterone during the three minutes after capture (Baugh et al., 2017a, 2013). A handful of other papers also report differences in aspects of the speed of GC responses between isogenic lines (Sadoul et al., 2015) or in relation to individual characteristics such as age and dominance (Sapolsky, 1993; Sapolsky and Altmann, 1991), food availability (Heath and Dufty, 1998), prior experience (Cockrem, 2013), or maternal condition (Weber et al., 2018). In addition to variation between individuals, there is ample evidence that the time required to reach maximum GC levels differs with life history stage (Wingfield et al., 1992), among populations (Addis et al., 2011; Zimmer et al., 2020b), and among species (Romero and Reed, 2005; Vitousek et al., 2018), although these studies typically interpret variation primarily or exclusively in terms of scope.

Despite this evidence that the speed of the GC response varies and suggestions that this variation might be an important target of selection, there has been little effort to assemble a complete conceptual framework for predicting when faster or slower GC responses would be favored at either an individual or population level. In contrast, a wide range of conceptual and mathematical models have explored the conditions under which the scope of the GC response is expected to be larger or smaller (Romero et al., 2009; Taborsky et al., 2020; e.g., Wingfield et al., 1998). These models have been applied to empirical data at both the between individual and among species levels (Bokony et al., 2009; Hau et al., 2010; Jessop et al., 2016, 2013; Schoenle et al., 2018; Vitousek et al., 2019).

In this paper, our goal was to first develop a set of hypotheses and predictions describing the conditions under which faster or slower GC responses should be favored. For this goal we borrowed heavily from existing frameworks for understanding variation in scope and translated these predictions to a set of hypotheses that might explain variation in speed of the GC response between individuals or populations (see below). We also evaluated support for predictions about how the speed and scope of GC responses covary between individuals and among species.

To evaluate evidence for these hypotheses, we used a database of corticosterone measurements in birds. The data available were more appropriate for testing differences in speed of GC regulation among species and we focus on those comparisons, but we emphasize that each of our hypotheses could also apply at the between individual level and that different patterns of covariation might occur at each level (Agrawal, 2020). Finally, we lay out recommendations and directions for future study in this area. Throughout the paper, we focus on the acute GC response because most empirical data include measurements of this aspect of the stress response, but many of the hypotheses and ideas developed here will apply equally well to other components of the integrated stress response that change rapidly after encountering a stressor. Measuring multiple aspects of the acute stress response to evaluate whether a faster GC response always results in faster downstream changes in phenotype will be a fruitful area for future study.

### Covariation in speed and scope

The speed and scope of endocrine responses could covary due to shared regulatory mechanisms, or as a result of selection operating simultaneously on both traits. Although phenotypic correlation does not necessarily equate to genetic correlation, absent or weak phenotypic correlation between these traits would suggest that they could be independently shaped by selection. Covariation between speed and scope is also important to understand because the particular patterns of covariation and relative amount of variation in each trait will have a strong effect on how well particular experimental designs can separately measure speed and scope (Taff, 2021). A recent optimality model by Luttbeg et al. (2021) revealed that slower GC responses lead to more similar baseline and stress-induced GC levels (i.e., a lower scope of response) when the increased lag time between encountering a stressor and responding appropriately elevates the likelihood of a mismatch between context and hormonal state. Here, we tested whether these predictions are supported at the between individual and among species levels. Specifically, we tested whether individuals and species that mount a faster GC stress response have lower baseline GCs and a larger GC scope (maximum - baseline). While Luttbeg et al.’s (2021) model considers corticosterone on a single scale, we evaluated covariation on both an absolute and relative (species centered) scale (see below and Box 1).

### The environmental and life history predictors of rapid GC responses

We predict that selection will favor faster GC stress responses in environments in which significant challenges are common - and in which the effects of those challenges could be ameliorated by rapid hormone-mediated plasticity. This overarching hypothesis is similar to the “supportive” hypothesis previously proposed to explain variation in baseline GCs and the scope of the acute stress response (Vitousek et al., 2019); however, we anticipate that the specific environmental and life history contexts that most strongly favor a rapid response versus a high scope response will differ. Because of the role of GCs in mediating thermoregulation through metabolic effects and the response to environmental challenges (Debonne et al., 2008; e.g., Jessop et al., 2016; Ruuskanen et al., 2021) we predict that: (1) faster GC responses will be favored in environments with greater thermal variability and/or unpredictability, and possibly also (2) in environments with greater variability or unpredictability in rainfall. We also predict that because smaller organisms generally have fewer energetic reserves, selection will favor (3) a more rapid GC stress response in smaller species. Similarly, when controlling for body size, we predict that (4) species with a higher metabolic rate (and thus higher total energetic demand) will mount faster GC stress responses. Note however that a positive covariation between metabolic rate and the speed of GC responses could also be a byproduct of the generally faster rate of biochemical processes that accompany high metabolic rates, rather than selection specifically favoring fast GC stress responses in these species.

Because mounting a GC stress response imposes a variety of costs, selection may also favor a muted GC stress response in contexts in which these costs are likely to be particularly damaging (the “protective” hypothesis: Vitousek et al. 2019). If a slower GC stress response reduces the likelihood that a response will be triggered inappropriately by challenges that cease before the onset of GC-mediated plasticity, or provides individuals with more time to evaluate the nature of a challenge before responding, then slower responses may be especially beneficial in some contexts (Luttbeg et al., 2021; Taff and Vitousek, 2016). We predict that because the acute GC stress response often impairs reproduction (e.g., Bokony et al., 2009; Sapolsky et al., 2000; Wingfield and Sapolsky, 2003), (5) organisms engaging in high value reproductive attempts (those with fewer lifetime opportunities to reproduce) will mount slower stress responses during breeding.

The nature of the challenges that organisms face are likely to affect the optimal speed of GC responses, in addition to their scope (e.g., Schoenle et al., 2018). When predation and other extrinsic threats are variable in frequency, and when the risk of these threats can be mitigated by GC-induced plasticity, then we predict more rapid responses will be favored in populations that encounter these threats more often. Because data on the frequency or nature of threats faced by individuals in the populations measured here are not available we were not able to test this prediction directly. However, we tested the related prediction that (6) shorter-lived species (which generally face more extrinsic threats) will mount faster GC responses. Note however that this same relationship could reflect selection favoring slower responses in longer-lived species, which may be more susceptible to accumulated phenotypic damage resulting from high GC levels (Schoenle et al., 2021; Vitousek et al., 2019).

## METHODS

### Database of corticosterone measurements

We used a database of corticosterone measurements taken from species studied by the Wingfield Lab between 1988 and 2005 (Wingfield et al., 2018, 1995, 1992). Most of these data have been published previously as parts of individual studies spanning the last several decades. Baseline and stress-induced corticosterone values for most species are also included in HormoneBase (Vitousek et al., 2018), but that database does not include data from each time point used here. The field and laboratory methods for these studies are similar across species and are described in detail in a number of previous papers (Wingfield et al., 1995, 1992).

For all species, individuals were captured and a blood sample was taken in under three minutes followed by a standard stress restraint protocol with samples taken at multiple time points after capture. Samples were stored on ice in the field until plasma and red blood cells were separated by centrifugation in the lab and corticosterone concentration was assayed by radioimmunoassay (Wingfield et al., 1995, 1992). No new data were collected in the present study. All sampling was approved by the appropriate agencies spanning a variety of institutions and locations.

Because we were interested in assessing variation in the speed of the corticosterone response, we restricted our analyses to species that had at least 5 individuals sampled for at least three different time points under 35 minutes after capture. For most species, samples were collected at <3 minutes, 5 minutes, 10 minutes, and 30 minutes. A few species had samples taken at 15 or 20 minutes in place of one of the other sampling times and a small number of samples were collected in between these standard times. The exact latency was recorded for all samples and often samples targeted to a specific time were actually collected a minute or two before or after the target time. We binned sampling times to determine species inclusion, but in all models we used the exact latency as a continuous predictor except in the case of some baseline samples (see explanation below). After filtering, our dataset included 60 species. Of these, 55 species also had at least 5 individuals sampled at a later time point (usually 60 minutes). Thus, most species in the dataset were sampled at five different time points during the hour after capture.

The database we used included information on mass, sampling date, and location of each individual. We matched these records with life history variables previously assembled in HormoneBase (as described in Johnson et al., 2018) to include average lifespan, number of clutches per year, age at maturity, and metabolic rate (Vitousek et al., 2018). Following Vitousek et al. (2019), we calculated the number of reproduction attempts as (average lifespan - age at maturity) x number of clutches per year. Previous analyses in the HormoneBase project used imputed metabolic rate and average lifespan from a phylogenetic reconstruction for species with missing data (Vitousek et al., 2019) using the R package phylopars (Bruggeman et al., 2009). We ran analyses both with and without imputed values. We report the analyses with imputed values but note any cases where results differed.

Finally, we also used data from a previous HormoneBase analysis (Vitousek et al., 2019) at a population level to match corticosterone records with the amount of variation in precipitation and temperature at each location. Briefly, intra-season variation in temperature and precipitation was calculated as the standard deviation of daily temperature from a 51-year time series of global climate in 0.5^*°*^ grids from the Climatic Research Unit (Harris et al., 2014) as described in Johnson et al. 2018. For these calculations, climate data were grouped into four three month intervals as follows: December-February, March-May, June-August, September-November (full details in Vitousek et al., 2019). Individual capture records were matched to the climate data for the location and time period that they occurred in. Species level data were calculated by averaging climate data across each individual record included.

### Characterizing the speed of the corticosterone response

The ideal way to separate variation in the speed of the corticosterone response from the absolute levels of corticosterone would be to fully estimate the response curve for each individual and species and compare all of the critical parameters and their correlations as suggested in Box 1. However, some of these parameters, such as the timing of the inflection point leading to a rapid increase after disturbance, would require many samples with fine temporal resolution and it is clear that the data available are insufficient to distinguish those differences. There is some evidence that individuals differ in the timing of initial activation (Baugh et al., 2017b, 2013) and it may be possible to measure these differences using approaches such as cannulated animals with repeated sampling or by measuring many individuals in a group or species at a mix of different time points rather than at standardized times (Taff, 2021), but our dataset is inappropriate for these questions. Instead, we focus our analyses on parameters that can plausibly be estimated given the data available, while recognizing that we cannot fully characterize variation in the functional shape of the response.

In contrast to inflection points, the slope of individual and species level increases and the absolute value of corticosterone at fixed time points can be reliably estimated with relatively few sampling points. We focused on estimating the absolute value of corticosterone at baseline along with the absolute or relative speed of increase in the first 15 minutes after disturbance. The absolute rate of increase during this time is likely a reasonable approximation of the maximum rate of increase (Box 1). To model the relative rate of increase, we centered and scaled all corticosterone measurements within each species. This allowed us to model species that might have widely different ranges for the absolute values of corticosterone on a similar scale, where the slope represents the maximum rate of increase measured in units of within species standard deviations. Modeling on this relative scale is a reasonable approximation for the time required to reach a given percentage of the species maximum value (Box 1). Individuals and species that come closer to reaching the species maximum value of corticosterone in the first 15 minutes after disturbance will have a faster speed (assessed by random slope estimate) on this relative scale.

For most species, limiting samples to the first 15 minutes included sampling time points at approximately 1, 5, and 10 minutes with a few species instead having a sampling point at 15 minutes. We chose to focus on samples taken in the first 15 minutes rather than all time points for several reasons. First, corticosterone values increased approximately linearly over this time period, making it possible to estimate slopes without requiring transformations or non-linear fits (Figure 2). While log transformation might have resulted in a linear increase over a longer time period, it would make it difficult to evaluate differences in the speed of response, because on a log scale the same absolute rate of increase would differ in slope depending on baseline corticosterone. We caution that studies interested in disentangling speed and scope will need to be very careful about the application and interpretation of transformations.

**Figure 2:**
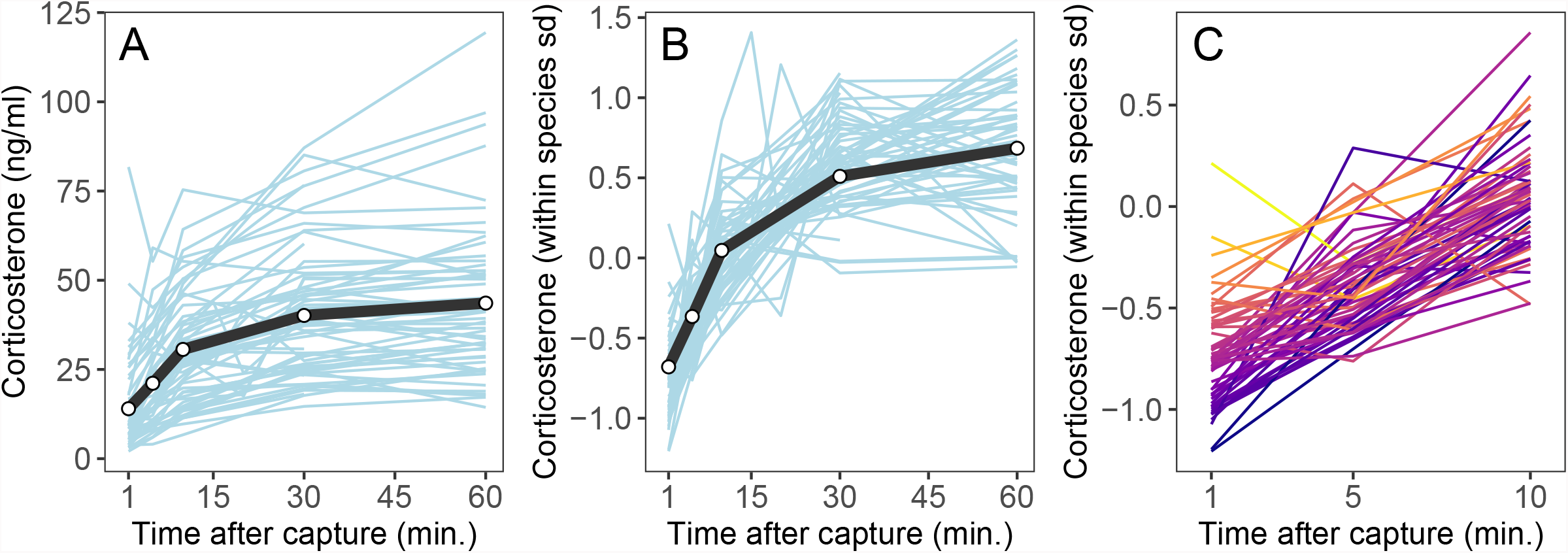
Comparative data from 60 species showing the overall absolute stress response based on raw data (A) and the same data with corticosterone standardized within species (B). Panel C reproduces panel B but is zoomed in to show differences in relative speed over the first 10 minutes more clearly. Each species is represented by a different line; in panel C, lines are colored by initial baseline values. Black line and points show the overall mean across all species at bins for under 3, 5, 10, 30, and 60 minutes. Note that the sample sizes for each species and for each bin within species vary considerably and this plot is meant as an illustration only. These plots show raw averages for each species and bin, but analyses are based on full data for each individual as described in the text.

Second, the maximum rate of corticosterone increase occurs in the first few minutes after a disturbance (Box 1) and this maximum rate should be best captured by the linear slope early in the response. Third, the slope over the entire sampling period may be related to maximum and baseline levels for purely mathematical reasons, especially when the exact time of individual and species maximum values are not estimable (e.g., if samples are only available at the exact same two time points, then slope will be mathematically identical to the scope, Figure 1). Finally, because we could not reliably estimate the exact time needed to reach maximum values with the available data, slopes using the full time course will include a mix of samples taken before and after the maximum and the offset of these samples from the maximum time may differ among individuals and species. When plotting raw data for each species, maximum values were always reached more than 15 minutes after disturbance (Figure 2) and using this time cutoff ensured that we were comparing species based on the slope of initial increase before reaching their maximum.

We used the full time course of samples to calculate an estimate for the scope of response for each individual (maximum - minimum at any time points). This scope was used in bivariate models exploring correlations between the speed and scope of the corticosterone response between individuals and among species, but was not included in the life history models.

### Considerations for estimating baseline corticosterone

In order to model correlations between the speed and absolute values of the corticosterone response we parameterized models so that the intercept corresponded to the approximate baseline level. This strategy allowed us to directly evaluate correlations between random slopes and intercepts themselves and between random estimates and life history covariates at both individual and species levels within the same models while appropriately accounting for the uncertainty in random effect estimation (Houslay and Wilson, 2017). Fitting models with this strategy required us to make decisions about baseline values that are somewhat subjective.

We needed to account for the fact that there is a lag between capture and the detection of higher circulating corticosterone (Wingfield et al., 1998). For birds, this lag time is often considered to be 3 minutes (Small et al., 2017), but the exact time is not known with precision and individuals or species could vary in the length of this time lag (Baugh et al., 2017b, 2013). As described above, we cannot estimate differences in time lags given our data.

Instead, we assumed a standard time lag of 2 minutes and counted any samples taken in under 2 minutes as baseline (set to a latency of 0 in our models). Using this approach, the intercept in our baseline models represented estimated corticosterone 2 minutes after capture. We performed a sensitivity analysis using a time lag of 1 minute or 3 minutes and all results were qualitatively similar. We acknowledge that this approach does not account for differences in time lags. If time lags vary systematically with speed (e.g., shorter time lag correlated with steeper initial slope), then some of the effects we detect could be associated with time lags rather than slope, but it is impossible to model those patterns with the data presently available.

### Data analysis

In all of our analyses, we modeled corticosterone changes after disturbance as within-individual reaction norms, drawing on the growing literature and methods that have been developed for this type of data from behavioral studies (e.g., Allegue et al., 2017; Hertel et al., 2020). A number of conceptual reviews have suggested that physiological stress responses should be considered as reaction norms (Hau et al., 2016; e.g., Taff and Vitousek, 2016; Wada and Sewall, 2014), but relatively few empirical studies have explicitly taken this approach to date (but see, Fürtbauer et al., 2015; Houslay et al., 2022) and we are not aware of any comparative papers that have modeled corticosterone responses as reaction norms.

We initially asked whether the speed of the acute corticosterone response (the random slopes) covaried with baseline corticosterone (the random intercepts) at both a within species and between species level. We addressed this question with two similar models using either absolute or relative corticosterone measurements taken in the first 15 minutes after sampling. These two models allowed us to evaluate the correlation of baseline corticosterone and the scope of the response with the speed of the absolute or relative corticosterone increase, respectively.

In each bivariate model, absolute or relative corticosterone was one response with latency after increase as a single fixed predictor. We modeled random slopes and intercepts for both individuals (to account for repeated samples in a series) and species (to account for multiple individuals per species). Scope was modeled as a second bivariate response including only the species level random effect structure to allow us to estimate correlations between scope and speed. For the species level random effect, we included the covariance matrix based on phylogeny to account for non-independence. To construct the covariance matrix we downloaded a resolved phylogeny from www.birdtree.org and pruned the tree to include only the species included in this study (Jetz et al., 2014, 2012).

Models were fit using the brms package in R, which passes models to Stan for markov-chain monte carlo sampling using the no-U-turn sampler (Bürkner, 2017; Carpenter et al., 2017). We generally used the default settings in brms for prior specification, warm up, and number of iterations with 4 chains per model and the default non-informative priors. All models were assessed visually using ShinyStan following the recommended diagnostics (e.g., checking for chain mixing, effective sample sizes, and Rhat) (Team, 2017). In a few cases, we increased the number of iterations to reach the effective sample sizes recommended by Stan. We interpret results as supported when the 95% confidence interval did not cross zero based on the full posterior distribution from fit models.

Next, we asked whether variation in the speed of the stress response was associated with life history variables at the species level by fitting one bivariate model corresponding to each hypothesis. We began with the exact same model specification described above with either absolute or relative corticosterone and one life history variable as the bivariate responses. The predictors for the corticosterone response variable were exactly as described above. For the life history variable, the only predictor was the random effect of species to account for the non-independence of life history variables measured from related species.

The life history variables used were (1) intra-season temperature variability, (2) intra-season precipitation variability, (3) log transformed mass, (4) metabolic rate corrected for log body size (residuals), (5) average lifespan, or (6) average lifetime reproductive attempts (reproductive value). For metabolic rate, lifespan, and reproductive attempts, we fit models with and without imputed values. For these three variables we had only a single measure for each species and it was therefore not possible to get reliable estimates of the amount of uncertainty in species level random effect estimates. We proceeded with analyses using the single available measure per species, but acknowledge that this approach likely underestimates the uncertainty in correlations derived based on these estimates.

After fitting models with this bivariate approach, we were able to directly assess the correlation between the life history variable of interest (random intercept accounting for phylogeny) and the species level random slope (speed) and intercept (baseline) while accounting for all uncertainty in random parameter estimation across both the individual and species levels. We report summaries of the full posterior distribution for these correlations.

Not all life history variables were available for all species and the sample sizes therefore vary between the models. Given the modest number of species included, and the fact that many of the life history measures we considered are likely correlated, we did not attempt to rank models and instead focus on cautious interpretation of each model separately while recognizing that we cannot separate the influence of each life history trait from the others. Because a larger analysis from the HormoneBase project has already explored similar analyses in relation to baseline and maximum glucocorticoids (Vitousek et al., 2018), we primarily focus on associations with speed.

For most models we used the full dataset, but the reproductive value hypothesis applies specifically to samples collected during the breeding season. Thus, for that model we restricted the dataset to individual samples collected during March to August for north temperate species and September to February for south temperate species. When samples were collected from populations located within 20 degrees of the equator, and from individuals whose breeding status was unknown, we considered them to be from the breeding season if the months of collection overlapped with the breeding season of that species.

## RESULTS

In total, our analysis included 7,074 corticosterone measurements from 1,750 individuals sampled from 60 different species. These species varied substantially in their absolute levels and rates of increase in circulating corticosterone (Figure 2A). When placed on a relative scale (centered and standardized within species) there was still variation in GC regulation, with some species coming closer to achieving their own maximum value (regardless of the absolute level) within the first 15 minutes after disturbance (Figure 2B). Even on the relative scale, species varied considerably in the rate of increase in the first few minutes after disturbance (Figure 2C).

### Covariation between speed and circulating corticosterone

When modeling the absolute values of corticosterone in the first 15 minutes after disturbance, the speed (rate of increase) was positively correlated with baseline concentrations both between individuals and among species. Similarly, the speed of this response was positively correlated with the total scope of the response both between individuals and among species (Table 1).

**Table 1.** Estimated correlation between the absolute or relative speed of increase in corticosterone and baseline levels or full scope of response. Correlations and 95% confidence intervals are based on the posterior distribution of two models described in the text. Correlations with confidence intervals that do not overlap zero are shown in bold.

| Comparison | Level | Absolute Speed of<br>Corticosterone Response | Relative Speed of<br>Corticosterone Response |
| --- | --- | --- | --- |
| Correlation of Speed<br>and Baseline | Between Individuals | **0.16 [0.01, 0.33]** | **0.31 [0.13, 0.52]** |
|  | Among Species | **0.44 [0.15, 0.70]** | **-0.75 [-0.90, -0.52]** |
| Correlation of Speed<br>and Scope | Between Individuals | **0.34 [0.31, 0.38]** | **0.26 [0.22, 0.30]** |
|  | Among Species | **0.93 [0.81, 0.99]** | 0.24 [-0.10, 0.56] |

In the model using corticosterone values standardized and centered within each species, the speed of the response was still positively correlated with baseline corticosterone between individuals. However, the correlation among species was reversed in this case with a strong negative correlation. The negative correlation among species indicates that species with high relative baseline values (i.e., baseline values closer to the overall mean corticosterone level for that species) had slower initial increases in relative corticosterone (i.e., a smaller percentage of the overall change in corticosterone was accomplished in the initial 15 minutes; Table 1).

The speed and scope of the corticosterone response were positively correlated between individuals in the relative corticosterone model. However, there was no correlation between speed and scope among species in the relative corticosterone model (Table 1).

### Life history traits and variation in speed

In models using the absolute values of corticosterone, there was no evidence for a correlation between any life history trait and the absolute rate of corticosterone increase over the first 15 minutes after disturbance (Figure 3). However, there was a trend for smaller species to have a faster absolute corticosterone increase (Figure 3; correlation of body size and absolute corticosterone speed = -0.40, 95% confidence interval = -0.79 to 0.03).

**Figure 3:**
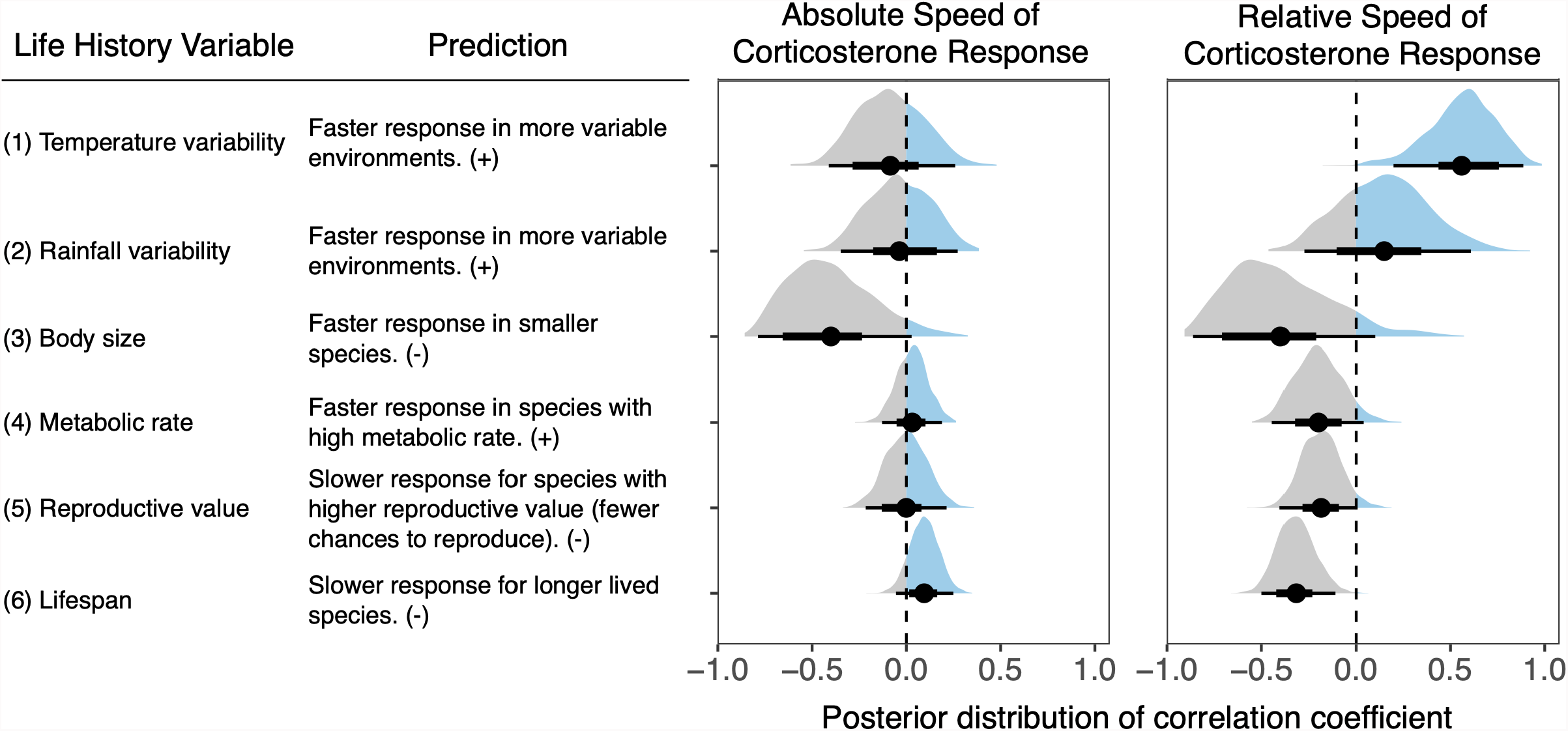
Posterior distribution of the correlation between the absolute or relative speed of the corticosterone response over the first 15 minutes after disturbance and each of the life history variables evaluated. The full posterior distribution is shown along with the mean (circle), the 66% confidence interval (thick line), and the 95% confidence interval (thin line). Each distribution is derived from a separate model fit as described in the text.

In models using the relative (species centered and standardized) values of corticosterone, there was strong support for a positive correlation between corticosterone speed and the amount of temperature variability (Figure 4A; correlation = 0.56, 95% CI = 0.20 to 0.89) and a negative correlation between speed and species average lifespan (Figure 4B; correlation = -0.32, 95% CI = -0.50 to -0.11). None of the other life history variables were clearly correlated with relative corticosterone speed, but there were trends for negative correlations between speed and body size (correlation = -0.40, 95% CI = -0.86 to 0.10), mass corrected metabolic rate (correlation = -0.20, 95% CI = -0.44, 0.04), and reproductive value (Figure 3; correlation = -0.19, 95% CI = -0.40 to 0.01).

**Figure 4:**
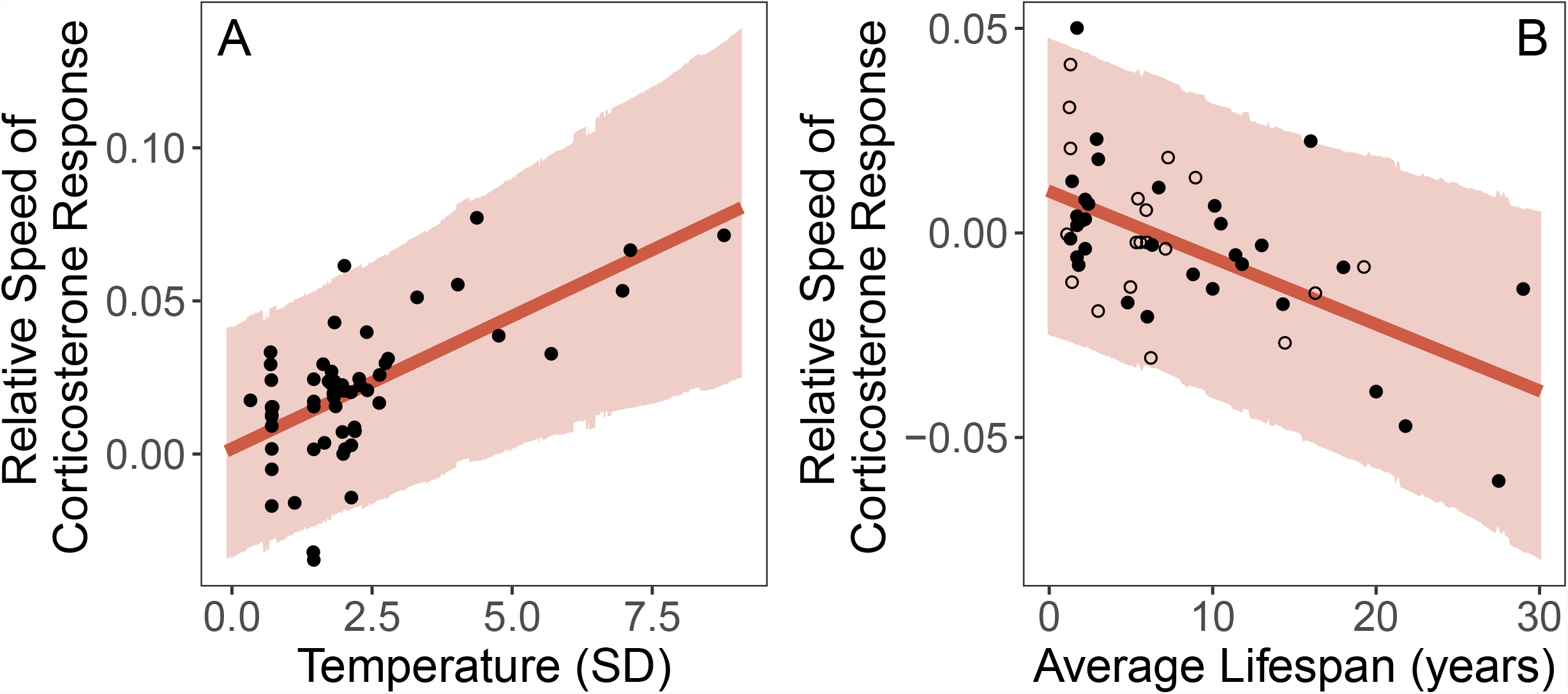
Model predicted relationship between the relative speed of the corticosterone response and temperature variability (A) or average lifespan (B). Points show the random effect estimates for each species. In panel B, species values that were inferred from a previous phylogenetic analysis are shown as open circles. In both panels, the line illustrates the mean correlation coefficient and shaded region shows the 95% confidence interval of the correlation from the full posterior distribution.

## DISCUSSION

While the factors shaping selection on the scope of GC responses have been well described in recent years, much less is known about whether variation in the speed of the GC response is also an important trait. Our results support the general idea that the speed of the acute GC response may be a target of selection both through its association with the scope of the GC response and via independent associations with environmental context or important life history characteristics. In particular, the relative speed of the response rather than absolute speed was associated with life history variables. At present, it is unclear under what conditions variation in speed or scope contribute more to fitness outcomes, largely because the available data in many published studies cannot distinguish between speed and scope. Nevertheless, our results suggest that the speed of the GC response, independent of scope, may play a role in determining how individuals and species cope with challenging environmental conditions.

The patterns of covariation that we found between the speed and scope of the acute GC response were largely similar to those predicted by the optimality model of Luttbeg et al. (2021). In absolute terms, there was a strong association between the scope of the GC response and the rate of increase during the initial 15 minutes after disturbance at both the between individual and among species levels. This pattern is consistent with the hypothesis that slower GC elevation will result in a smaller scope with more similar baseline and maximal GC levels to minimize the amount of time spent in a mismatched (suboptimal) phenotype (Luttbeg et al., 2021). In contrast with the strong association between scope and absolute speed, scope and the relative speed of the corticosterone response were not correlated among species. While individuals with larger scopes were still faster in this relative measure, there was no overall species-level association. The fact that the absolute rate of increase in corticosterone and the relative increase show different patterns suggests that - at least at the interspecific level - relative speed and scope could vary somewhat independently and may be subject to different selective pressures. More studies are needed that can separately measure speed and scope to assess the relative importance and amount of variation in these two traits, especially at the within-species level (Taff, 2021).

We also found that the speed of the acute GC response was positively correlated with baseline GC levels between individuals on both the absolute and relative scale. This result is somewhat surprising given the prediction from Luttbeg et al.’s optimality model (2021) that baseline GCs and speed should be negatively correlated. It is unclear what caused this discrepancy, but one possibility is that mechanistic constraints produce this association. Alternatively, it is possible that the pattern is caused by a tight correlation between the time lag to begin increasing and the speed of increase. Individuals in some species are known to differ in their timing of the onset of GC increase (Baugh et al., 2017b, 2013). Given the structure of our data collection and analysis, a shorter time lag to increase should result in a higher baseline estimate; if those individuals also have a faster rate of increase it could produce the between-individual pattern we observed even in the absence of a correlation with true baseline values. We found a similarly positive correlation between baseline and speed at the among species level for absolute corticosterone, but this correlation was reversed on the relative scale where species with relatively high baseline values had slower relative rates of GC increase. This among species pattern does match the predictions of an optimality model (Luttbeg et al., 2021) and we suggest that among species comparisons on a relative scale may provide a more accurate description of differences in the speed of downstream consequences of GC elevation (Box 1).

Among the environmental and life history factors tested, the strongest predictor of the relative speed of GC responses in birds was thermal variability. Species inhabiting environments with more intra-season variation in temperature mounted faster relative GC responses. This is consistent with the hypothesis that the ability to mount a rapid GC response to thermal challenges may be favored in highly variable environments and suggests a “supportive” effect of selection. In contrast, variation in precipitation did not predict the speed of GC responses in birds, though the trend was in the predicted direction. A previous analysis found that variation in both temperature and precipitation positively predicted baseline GC levels across vertebrates; this was interpreted as reflecting the role of baseline GCs in helping organisms to prepare for and cope with energetically demanding environments (Vitousek et al., 2019). We suggest that the different patterns seen here in the relationships between the speed of GC responses and variation in temperature and precipitation could reflect a difference in the timescale of the threat posed by these challenges: while extreme temperatures can represent an immediate threat to survival – for which it can be important to respond rapidly – variation in precipitation likely challenges birds over longer timescales (days to weeks). Thus, the relative benefit of responding rapidly to challenges may be greater in more thermally variable environments than in those that vary in precipitation. Alternatively, the difference might simply reflect the fact that analyses of scope have used a larger number of species with more variation in environment.

Shorter-lived species also mounted faster GC responses, when speed was measured on the relative scale. This pattern could reflect selection favoring more rapid stressor-induced plasticity in populations that face more extrinsic challenges (in accordance with the “supportive” hypothesis). However, the same relationship could also result from selection favoring slower responses in longer-lived species, who may be more at risk of accumulated phenotypic damage from elevated GC levels (“protective” hypothesis).

Contrary to our predictions, we did not find a significant relationship between lifetime reproductive attempts and any of the measures of the speed of the GC response during the breeding season. Thus, we found no clear support for the prediction that birds engaging in more valuable reproductive attempts (those with fewer lifetime reproductive opportunities) reduce the likelihood of GC-induced reproductive impairment by responding more slowly to threats. We did, however, find a trend in the predicted direction and it is possible that a larger sample of species would support this prediction. It is also important to note that the various life history measures that we assessed were tightly correlated in this data set; species with greater longevity also had more lifetime opportunities to reproduce. Thus, while longevity is clearly a stronger predictor of the speed of GC responses than reproductive value in this dataset, the non-independence of these measures prevent us from determining the extent to which reproductive value may independently predict the speed of GC responses.

Neither body mass nor metabolic rate were associated with the relative speed of GC responses in birds, though there was a trend for a negative correlation with both. Previous analyses in birds and across vertebrates found that smaller species have higher baseline GCs (Bokony et al., 2009; Hau et al., 2010; Vitousek et al., 2019) but that size is unrelated to stress-induced GCs (Bokony et al., 2009; Vitousek et al., 2019; but see Hau et al., 2010). These findings suggest that body size alone may not predict whether a faster or slower GC response is optimal, although both absolute and relative speed tended to be negatively correlated with body size as we predicted. Despite widespread predictions that metabolic rate is a major driver of variation in GC release and clearance, metabolic rate appears to generally be a rather poor predictor of variation in GC levels across species. Mass-specific metabolic rate is not related to baseline GC levels within birds (Francis et al., 2018) or across vertebrates (Vitousek et al., 2019; but see Haase et al., 2016 in mammals). Birds with higher mass-specific metabolic rates do have higher stress-induced GC levels (Francis et al., 2018), but this pattern is not present over larger taxonomic scales (Vitousek et al., 2019). While there was a trend suggesting a possible negative association between relative speed and metabolic rate, this pattern was the opposite of what we had predicted. The lack of a relationship between metabolic rate and the speed of GC responses seen here underscores that the speed of endocrine responses – like other GC regulatory traits – can evolve independently of metabolic rate. It also suggests that total energetic demand is not a strong predictor of the optimal speed of GC responses.

Taken together these findings suggest that selection favors rapid GC responses in organisms facing frequent major challenges – consistent with the “supportive” role of GCs. In contrast, there was little definitive support for the idea that slower GC responses may help to protect organisms from the costs of over responding – and thus be favored in contexts in which the costs of mounting a GC response are particularly high (the “protective” hypothesis). Note however that as described above, the observed relationship between longevity and the speed of GC responses could reflect selection favoring either “supportive” or “protective” roles. A recent phylogenetic comparative analysis found a similar overall pattern for baseline corticosterone: across vertebrates, baseline GCs are higher in populations and species in more challenging environments, consistent with the “supportive” hypothesis (Vitousek et al., 2019), and also with “permissive” and “preparatory” roles for baseline GCs in helping organisms to respond to stressors (Sapolsky et al., 2000). Variation in peak stress-induced corticosterone was instead best explained by selection favoring reduced costs (the “protective” hypothesis). Understanding how the speed of GC responses is related to the frequency of challenges has important implications for predicting how species will respond to climate changes that result in increased frequency, duration, and intensity of extreme weather events.

The initial speed of the stress response might also play an important role in activating the rapid effects of steroid hormones. In general the mechanisms of action by non-genomic receptors are not well understood, but the perspectives presented in this paper may direct hypotheses and experimental approaches relevant to environmental context and speed of the acute stress response. Over several decades, evidence has been growing that steroid hormones can have very rapid effects, within minutes, which are not compatible with binding to genomic receptors. The latter act as gene transcription factors requiring hours for full response (e.g., Balthazart, 2021). Rapid effects of glucocorticoids in mammals, birds, and amphibians have been attributed to non-genomic receptors, possibly in cell membranes, that generate behavioral and physiological responses to environmental perturbations (Panettieri et al., 2019). Such effects include increased aggression in rats (Mikics et al., 2004), altered cell signaling (Haller et al., 2008), locomotion, anxiety and general behavior in response to an environmental challenge (Makara and Haller, 2001; Mikics et al., 2005). Non-genomic receptors for GCs appear to be associated with membranes in mammals (Tasker et al., 2005) and in amphibians these membrane receptors in the central nervous system interact with G-proteins, further suggesting non-genomic actions (Moore and Orchinik, 1994). In a songbird, non-invasive treatment with corticosterone (via ingestion of a mealworm injected with the steroid hormone) increased plasma levels of corticosterone and perch-hopping activity within 15 minutes (Breuner et al., 1998). It also appears that this rapid effect on activity is evident in birds held on spring-like long days and not manifest in birds held on winter-like short days (Breuner and Wingfield, 2000). Considering the initial speed of the acute stress response may lead to new insights into the cellular mechanisms by which more rapid GC responses allow for more effective avoidance or tolerance of stressors.

One limitation of this study is that we were only able to test life history related hypotheses at the among species level. There is evidence that variation in the scope of the GC response is related to life history traits or performance among species (Bokony et al., 2009; Hau et al., 2010; Jessop et al., 2013; Vitousek et al., 2019) and among individuals within a species (Breuner et al., 2008; Ouyang et al., 2011; Schoenle et al., 2021; Vitousek et al., 2014). Similar patterns may apply to speed, but few studies address speed at the within species or within individual level (but see Baugh et al., 2013) and simulations demonstrate that separately measuring speed and scope at these levels will be challenging (Taff, 2021). Moreover, while there is appreciation for the way that GC regulation varies across multiple levels (Hau et al., 2016), there is no guarantee that associations found at one level will apply at other levels (Agrawal, 2020). For example, here we failed to find a relationship between speed and average reproductive attempts. However, the species in our dataset varied enormously in lifespan and this variation may have masked the importance of variation in reproductive value between more closely related species. It is entirely plausible that a more narrowly focused analysis (e.g., between populations of the same species along a latitudinal gradient) would support the reproductive value hypothesis. Studies of both speed and scope would benefit from a focus on developing frameworks that explicitly make level-specific predictions (Agrawal, 2020; Hau et al., 2016).

We focused here on only the initial rapid increase in GCs after a stressor, but there are other timing related elements of the GC response that could be considered variation in speed (e.g., time spent at maximum, maximum rate of negative feedback, speed of steroid clearance, time to return to baseline levels; Box 1). Several recent papers have demonstrated that variation in the strength of negative feedback is an important predictor of performance (Romero and Wikelski, 2010; Taff et al., 2018; Zimmer et al., 2019). Interestingly, these results are sometimes interpreted as demonstrating variation in the speed of negative feedback even though measures are only taken at two time points, making it difficult to separate the scope and speed of negative feedback. Moreover, the speed of GC regulation represents only a single component of speed in the more general stress response (Romero and Gormally, 2019). There has been increasing recognition in recent years that GC regulation alone is insufficient to understand variation in the stress response, because a greater GC response does not necessarily indicate a greater response in a variety of important downstream physiological or behavioral traits (Gormally et al., 2020; Neuman-Lee et al., 2020; Romero and Gormally, 2019). While these studies have generally focused on variation in scope, the same arguments apply to understanding variation in speed. For example, different target cells or tissues themselves might differ in both the speed and scope of their response to an increase in circulating GCs. A more complete understanding of speed will require identifying the entire functional shape of acute GC responses and linking that response to downstream consequences.

Our results also suggest that it will be important to consider differences between speed on an absolute or relative scale. One interpretation of the patterns we found is that absolute differences in corticosterone scope are so large between species that they mask the importance of differences in speed. By standardizing each species on a similar scale, we were able to see that the speed with which species approach their species specific maximum value is associated with life history measures and varies somewhat independently from scope. If other aspects of the stress response system (e.g., receptors) co-evolve closely with variation in absolute corticosterone levels, then relative differences in speed (i.e., the shape of the response rather than magnitude) may be more relevant when considering the speed of downstream effects from mounting a GC response.

To some extent, there has been a growing appreciation for the need to understand flexibility in the shape of GC responses, even when speed and scope are not explicitly identified as potentially separate traits of interest. The recent emphasis on within-individual reaction norm approaches for studying variation in GC regulation (speed, scope, or the entire functional shape of responses) is an exciting development in this field (Hau et al., 2016; Taff and Vitousek, 2016; Wada and Sewall, 2014). However, we caution that these tools are still limited in many cases by available data and simulations demonstrate that creative study designs may be required to separately assess variation in speed and scope (Taff, 2021). Technical advances that allow for continual monitoring of GCs during an entire acute response under relatively natural conditions would be a huge step forward for this field. Regardless of the limitations, both the speed and scope of the acute GC response are clearly associated with important life history traits. Understanding how speed and scope covary or the conditions under which one or the other trait is a more important determinant of fitness may help to predict why some individuals and populations are able to survive in challenging conditions when others fail.

## ACKNOWLEDGMENTS

We are grateful to Patrick Kelley for compiling the endocrine data, and to the many people in the Wingfield Lab who helped to collect and assay plasma samples. David Westneat and three anonymous reviewers provided helpful comments on an earlier version of this study. We also thank the HormoneBase Consortium and Jennifer Uehling for compiling the life history, metabolic, and environmental data. CCT and MNV were supported by NSF-IOS 1457251 and 2128337, and DARPA D17AP00033.

## COMPETING INTERESTS

The authors declare no competing interests.

### Box 1

Defining and measuring the speed of acute stress responses

Conceptually, variation in the speed of the acute stress response is reflected by how quickly organisms can change their circulating glucocorticoid levels as conditions change (Taff & Vitousek, 2016). However, translating this broad definition to specific measurements reveals that there are several different aspects of the stress response that could be considered as representing variation in the speed of the acute glucocorticoid response and some of these cannot be easily separated from variation in the scope of the stress response, where scope is defined as the difference between baseline and maximum glucocorticoid concentrations (Figure 5).

**Figure 5:**
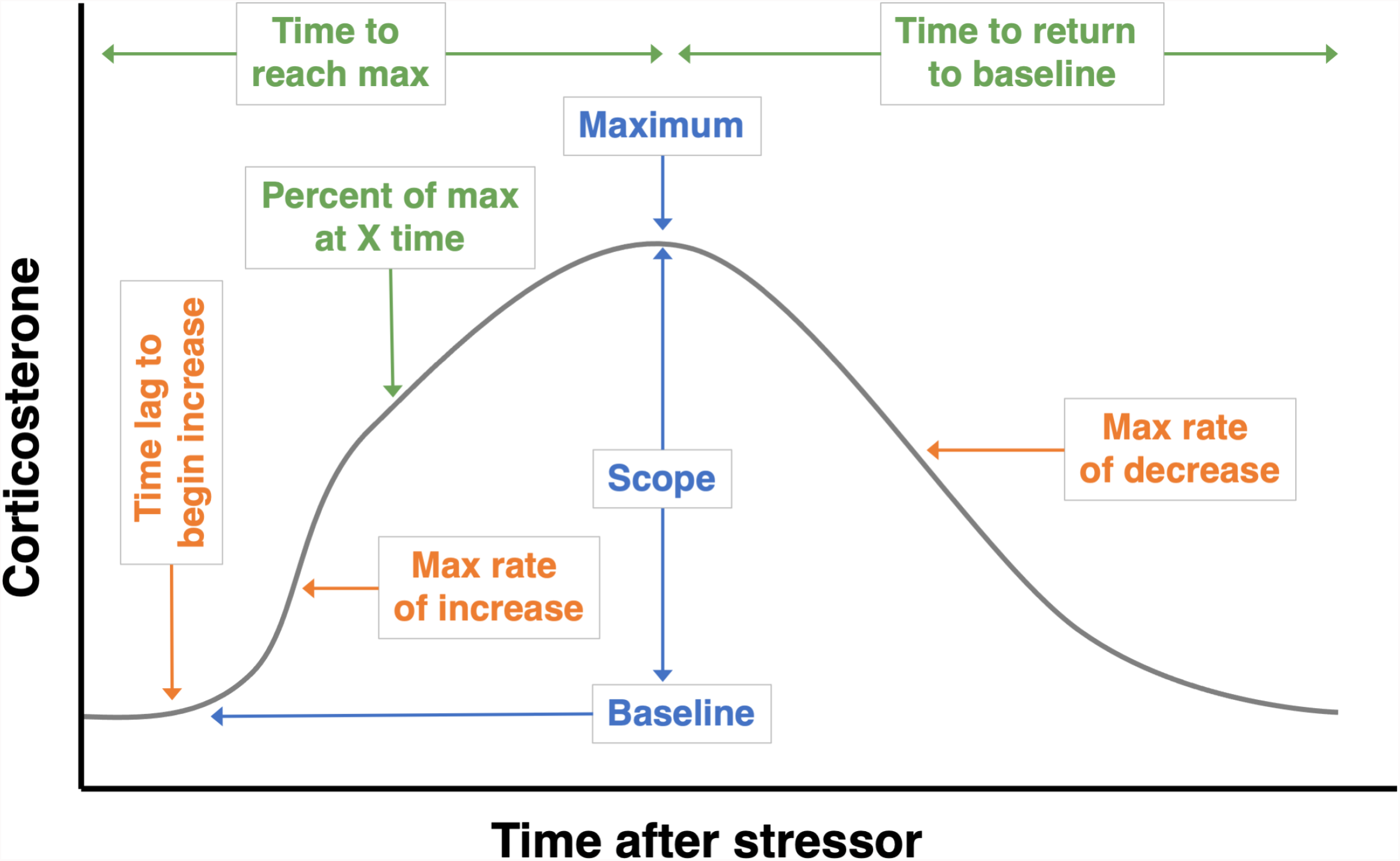
Possible measures that could be derived from a single continuous acute glucocorticoid response (gray curve). Traditional measures of absolute glucocorticoid levels or change are shown in blue (baseline, maximum, scope). Measures that directly reflect variation in speed are shown in orange (time lag to increase, maximum rate of increase and decrease). Three other potential measures are shown in green that, by definition, reflect a combination of speed and absolute glucocorticoids (percent of maximum or response at time X, time to reach maximum, time to return to baseline).

In theory, individuals or groups could vary independently in each of these aspects of the speed of acute responses, though in practice it may be common to find strong covariation between some components. It is worth noting that some of these measures are inextricably linked, such that speed and scope will both contribute to variation. For example, for any given maximum rate of increase, individuals with a higher maximum glucocorticoid value will, necessarily, require a longer time to reach their maximum. All of these measures are summaries of the variation reflected in the full response curve and which summary is the most informative may vary with the biology and patterns of variation present in any given study. Because species and individuals often vary enormously in the magnitude of the GC response, it may be more useful in many cases to make comparisons of speed on a relative scale (Figure 6).

**Figure 6:**
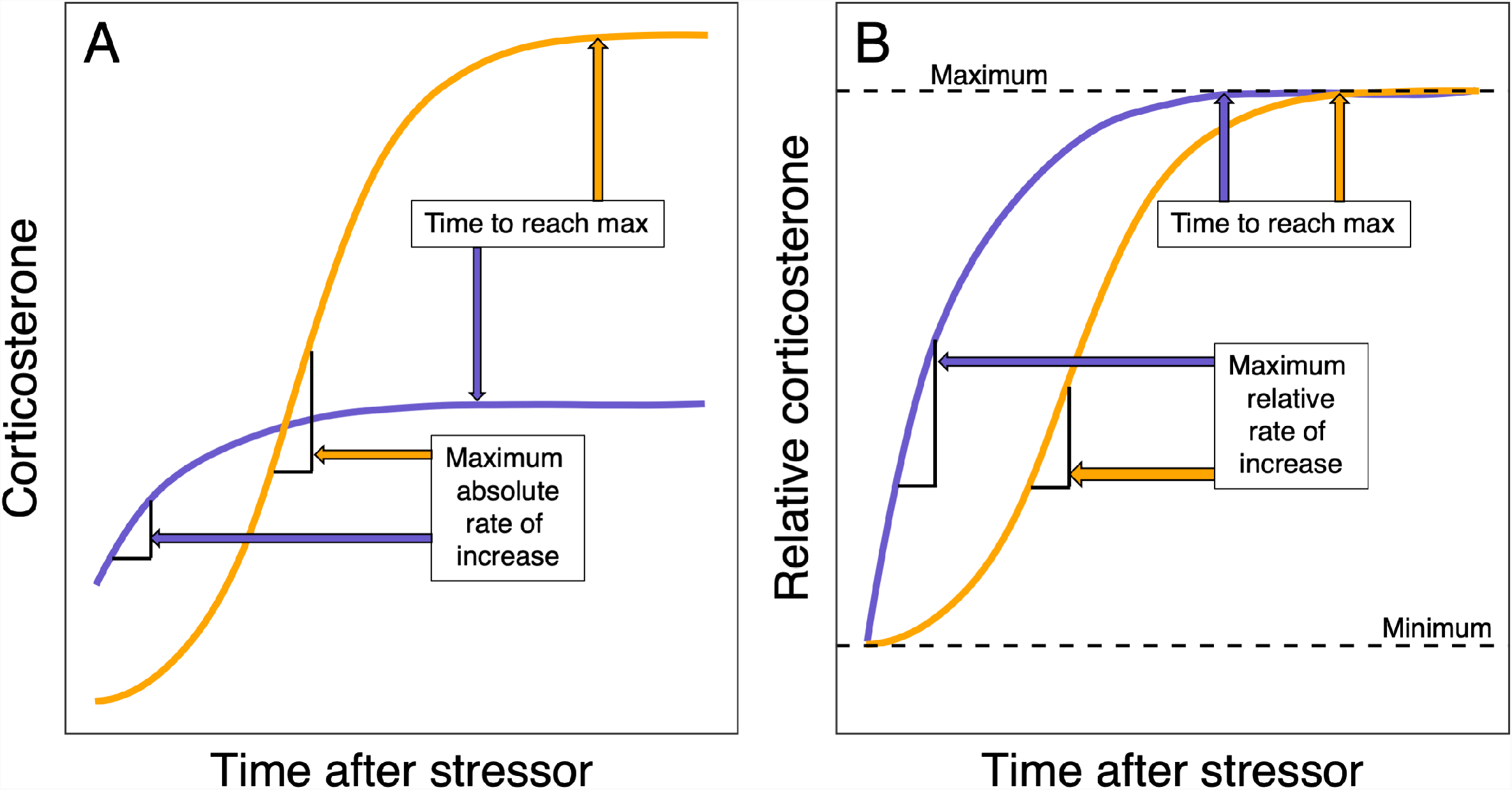
Comparison of two hypothetical corticosterone response curves that differ in both speed and scope plotted on an absolute (A) or relative (B) scale. Each curve could represent an individual or a composite group or species response. The time to reach maximum is unchanged on either scale, but the maximum rate of increase is strongly influenced by the overall scope of response such that the ranking of speed measured by rate of increase is reversed on the two scales. On the relative scale, the maximum rate of increase is likely to be a good predictor of time to reach maximum, but it may not be correlated with time to reach maximum on the absolute scale if there are large differences in absolute magnitude of the response. For simplicity, we do not illustrate downregulation here.

#### Time lag to begin increase

The time after a stressor is encountered to begin upregulation of circulating glucocorticoids.

#### Maximum rate of increase

The maximum rate of change of circulating glucocorticoids during an acute response. Will most likely be achieved during the early minutes of a stress response.

#### Maximum rate of decrease

The maximum rate of change of circulating glucocorticoids during the negative feedback phase after an acute response.

#### Time to reach maximum

The total amount of time from encountering a stressor to reaching the maximum circulating glucocorticoid level. Because this measure depends directly on the rate of change, maximum value, and baseline value, it will likely represent a combination of variation in speed and scope under most conditions.

#### Time to return to baseline

The total amount of time from reaching maximum glucocorticoid levels to return to baseline levels via negative feedback and clearance. This measure will likely represent a combination of variation in speed and scope.

#### Time to reach X percent of maximum

The amount of time taken from encountering a stressor to reaching a certain percentage of the maximum value. For example, species could be compared in how long it takes to reach 50% of their maximum value. As with the time to reach maximum, this will most often reflect a combination of speed and scope.

#### Time to reach X percent of scope

The amount of time taken from encountering a stressor to reaching a certain percentage of the acute glucocorticoid response (maximum - baseline values). This may differ from the percent of maximum because individuals or species that maintain high baseline glucocorticoids will start a response at a higher percentage of their maximum value. As with the other time measures above, this will most often reflect a combination of speed and scope.

#### Absolute vs. relative scales

Because most empirical studies will have incomplete data sampled at only a few time points, comparisons on a relative rather than absolute scale may be better at detecting some differences in the speed of the corticosterone response (Figure 6). In particular, comparisons of the time to reach maximum or percent of maximum reached by a certain time will often be easier to detect on a relative scale because these differences could be masked by large between individual or among species differences in overall magnitude of the response. Conceptually, comparison on a relative scale is similar to techniques such as Procrustes analysis or landmark based morphometrics used in morphological comparisons (e.g., Albertson and Kocher, 2001) or dynamic time warping analysis of similarity between two time series measurements used in kinematic or sound comparisons (e.g., Keen et al., 2021). This approach allows some aspects of speed differences to be compared directly even when there are large differences in the absolute levels of corticosterone between individuals, groups, or species.

